# Genome reconstruction and characterisation of extensively drug-resistant bacterial pathogens through direct metagenomic sequencing of human faeces

**DOI:** 10.1101/153874

**Authors:** Andre Mu, Jason C. Kwong, Nicole S. Isles, Anders Gonçalves da Silva, Mark B. Schultz, Susan A. Ballard, Glen P. Carter, Deborah A. Williamson, Torsten Seemann, Timothy P. Stinear, Benjamin P. Howden

## Abstract

Whole-genome sequencing of microbial pathogens is revolutionising modern approaches to outbreaks of infectious diseases and is reliant upon organism culture. Culture-independent methods have shown promise in identifying pathogens, but high level reconstruction of microbial genomes from microbiologically complex samples for more in-depth analyses remains a challenge. Here, using metagenomic sequencing of a human faecal sample and analysis by tetranucleotide frequency profiling projected onto emergent self-organising maps, we were able to reconstruct the underlying populations of two extensively-drug resistant pathogens, *Klebsiella pneumoniae* carbapenemase (KPC)-producing *Klebsiella pneumoniae* and vancomycin-resistant *Enterococcus faecium.* From these genomes, we were able to ascertain molecular typing results, such as MLST, and identify highly discriminatory mutations in the metagenome to distinguish closely related strains. These proof-of-principle results demonstrate the utility of clinical sample metagenomics to recover sequences of important drug-resistant bacteria and application of the approach in outbreak investigations, independent of the need to culture the organisms.

## Introduction

Clinical and public health microbiology is undergoing a major transformation driven largely by high-throughput microbial genome sequencing. The application of microbial genomics in these areas has been well described, including use for high resolution microbial characterization, source and transmission tracking for nosocomial and community pathogens, and antimicrobial resistance detection and prediction ^1^^-^^4^. Clinical and public health genomics, however, currently relies on routine culture-based assays to isolate pathogens of interest prior to whole genome sequencing, which presents inherent biases in analyses, and does not allow characterization of pathogens which are unculturable, below the level of culture detection, or unsuspected in a clinical sample ^5^^-^^9^.

Genomics-based approaches that overcome these difficulties by direct characterization of pathogens from clinical samples would be a major advance in clinical and public health microbiology. Metagenomics approaches can complement culture-based techniques for typing, and detecting, antimicrobial resistant (AMR) genes, and SNP variants ^10,11^. Metagenomics allows for the sequencing of whole community genomic material extracted directly from clinical samples, such as faeces, blood, cerebrospinal fluid, sputum, and bronchoalveolar lavage fluid ^12^,^13^. Current literature on public health metagenomics is largely based on interrogating the metagenome at ‘first-order’ level analyses; that is, either, characterizing bacterial biodiversity at the 16S rRNA gene level, identifying the functional profile of the microbial community, or alignment-based reference analyses for recovery of genomes ^12^. However, the challenge still remains in applying alignment-free analyses of metagenomic data to obtain strain-level resolution that might help understand transmission of pathogens in a clinical setting. Recent metagenomics advances in the field of environmental microbiology and ecology may provide potential solutions here, including techniques that bin contigs based on their tetranucleotide frequency profiles (c.f.,^14^^-^^17^).

In this proof-of-principle study, we used whole community metagenomics and pathogen genome reconstruction to interrogate the metagenome of a patient colonised with an extensively drug-resistant pathogen KPC-producing *K. pneumoniae*. Here we demonstrate that faeces metagenomics not only identified detailed SNP information to distinguish clonal *K. pneumoniae* isolates, but also uncovered unsuspected colonization with vancomycin-resistant *Enterococcus faecium* (VREfm), another high-risk antimicrobial resistant pathogen. Furthermore, we investigated whether metagenomic analysis could be used to characterise the resistance-harbouring genomes of a patient with long-term carriage of KPC-producing Klebsiella pneumoniae, and link that patient to a local transmission network, independently of the need for bacterial culture.

## Materials and Methods

### Epidemiological context

One faecal sample collected from a patient (Patient A) with known KPC-producing *K. pneumoniae* colonisation underwent whole community metagenomics. Patient A was a resident of an aged care facility and did not report any recent travel in the context of his comorbidities and frailty, but had been an inpatient at a tertiary hospital with a known outbreak of KPC-producing *K. pneumoniae* two months prior to sample collection.

#### Whole community genomic DNA extraction and high throughput metagenomic sequencing

Whole community gDNA was extracted from 0.2 grams of Patient faeces (herein referred to as AUSMDU00008155) using the QiaAMP Stool kit following manufacturer’s protocol with a preprocessing step of mechanical lysis (Bertin Technologies precellys 24). MP Biomedicals’ Lysing Matrix B 2-ml tubes containing 0.1 mm silica beads were used for two 40 second cycles of mechanical lysis (Bertin Technologies precellyis 24) at 6000x units with a 60 second rest on ice in between. Genomic DNA from the faeces, and a no-template control, were processed for sequencing using the Nextera XT kit on the Illumina MiSeq machine (V3, 600 cycles) (Illumina Inc, San Diego, US) following a modified manufacturer’s protocol. The following modifications were included: a 1% (v/v) spike-in ratio of PhiX, denatured DNA was diluted to a final concentration of 14.25 pM with pre-chilled HT1 buffer, and Tris-Cl 10 mM 0.1% Tween 20 was substituted with Qiagen’s EB solution to dilute sequencing libraries and PhiX throughout the protocol.

#### Culture-dependent whole genome sequencing

Concurrently, 14 individual colonies were picked at random from the same sample plated on Brilliance™ CRE selective media (Thermo Fisher Scientific, Waltham, US), with each colony undergoing whole-genome sequencing on the Illumina NextSeq 500 (Illumina Inc, San Diego, US). Two KPC isolates (herein referred to as AUSMDU00008118 and AUSMDU00008119) taken two days apart from a different patient that had previously undergone long-read sequencing to investigate the plasmid dynamics within an outbreak of KPC-producing *Klebsiella pneumoniae* were used as reference genomes for the analysis. DNA extraction, size selection, and sequencing on the Pacific Biosciences RS II (Pacific Biosciences, Menlo Park, US) were performed as previously described ^18^. Genomic DNA from these isolates was also sequenced on the Illumina NextSeq 500 for polishing to produce high quality closed genomes.

### Bioinformatic analyses

#### Metagenomic sequence data processing

Metagenomic data from AUSMDU00008155 were processed prior to analysis with Trimmomatic (v0.33) ^19^ for quality control and to remove adaptor sequences, PhiX contamination, and trace contaminants from Illumina preparation kits. Paired-end reads were merged and assembled using Iterative de Bruijn Graph De Novo Assembler for Uneven sequencing Depth (IDBA-UD) ^20^ compiled for long reads (i.e., 651 bp). Further quality control included removing host-derived gDNA using DeconSeq and the Human Genome Reference Sequence (build 38; GCA_000001405.22) prior to downstream analyses.

#### Metagenomic binning

To reconstruct isolate genomes from the gut microbial community, an emergent self-organizing map (ESOM) was used. Tetranucleotide frequencies were calculated for the assembled contigs using Perl scripts developed by Dick et al., (2009) ^16^ in preparation for analysis using ESOMs. The primary map structure was determined using in silico fragmented (> 5kb) contigs; while contigs between 2.5kb and 5kb in length were projected onto the ESOM using their tetranucleotide frequency profiles. Genomic binning was analysed using Databionic ESOM Tool with default settings except K-Batch training algorithm in 200x400 windows, a starting value of 50 for the radius, and data points were normalized by RobustZT transformation. Contigs with a native size smaller than 2.5kb were removed from analyses. Reference genomes were included in the analysis to guide identification of “binned” genomes, and validate completeness of genome recovery.

#### Detection of antimicrobial resistance genes

Assembled metagenomic contigs were screened for the presence of antimicrobial and virulence genes, including carbapenemases (*blaKPC*), using ABRicate (https://github.com/tseemann/abricate). Briefly, ABRicate detects acquired resistance genes using BLAST+ against the Resfinder database (Center for Genomic Epidemiology, University of Denmark^21^).

#### *In silico* molecular typing and detection of antimicrobial resistance genes

Assembled metagenomic contigs were screened for multi-locus sequence typing (MLST) scheme alleles using mlst (https://github.com/tseemann/mlst), an in-house tool that uses a BLAST algorithm ^22^ to search against the entire reference database of MLST profiles (downloaded from https://pubmlst.org). In addition, acquired antimicrobial resistance genes, including carbapenemases (blaKPC), were detected using another custom BLAST tool, ABRicate (https://github.com/tseemann/abricate), to search against the ResFinder v2.1 database ^21^

#### Reconstructing 16S rRNA gene squences

Near-complete 16S rRNA gene sequences were reconstructed from Patient B unassembled short read metagenomic data (post removal of host-derived gDNA) using the Expectation Maximization Iterative Reconstruction of Genes from the Environment (EMIRGE) program ^23^. The following parameters were incorporated: the SILVA Small Subunit database was employed as a training reference set, length of reads of 151, insert size of 683, standard deviation of 68, and a phred score of 33 were selected to compute over 80 iterations. Reconstructed 16S rRNA genes were queried against the Ribosomal Database Project using BLAST. A k-mer based approach, using Kraken ^24^, classified unassembled read data to support EMIRGE results.

#### Reference genome assembly to validate metagenomic bins

Reference genomes were assembled using Canu v1.5^25^, trimmed (https://github.com/tseemann/berokka) and circularized. Illumina short read data from the same gDNA sample were used to correct and polish the draft PacBio genomes using Pilon v1.22 ^26^ and Snippy v3.2 (https://github.com/tseemann/snippy). Further assembly of unmapped short read data (i.e., Illumina reads that did not match chromosomal or larger plasmid DNA from PacBio-derived data) using SPAdes v3.10.1 ^27^) was used to detect the presence smaller plasmids potentially missed through DNA size selection. Prokka v1.11 ^28^ was used to predict CDS regions and annotate the assembled genomes. Further characterisation of reference genomes including multi-locus sequence typing and antimicrobial resistance gene detection was performed in silico using the in-house developed tools, mlst and ABRicate as described above.

#### Transmission cluster inference

To determine the most likely transmission cluster source for Patient A, previously sequenced PacBio reference genomes from three local transmission clusters were assembled using the methods described above, and used to build a custom Kraken database. The local transmission clusters were defined through phylogenetic analysis of a maximum likelihood tree described in Kwong *et al.,* (*in prep*). Whole-community metagenomic sequencing reads were analysed in Kraken v0.10.5-beta ^24^ using the custom database to identify the most closely related reference genome.

## Results

### *In silico* typing and detection of AMR genes

Analysis of the assembled metagenomic contigs from Patient A identified the presence of two complete mlst profiles – ST258 *K. pneumoniae* and ST555 *E. faecium*. Table 1 highlights the resistance AMR genes detected with 100% coverage. The gene encoding resistance to carbapenem, *bla_KPC_*, was detected at 100% coverage and nucleotide identity. The following genes, with percentage coverage and nucleotide identity given in parentheses, were also recovered from metagenomic data: *vanR-B* (100, 99.2), *vanS-B* (100, 99.6), *vanY-B* (100, 100), *vanW-B* (100, 97.6), *vanH-B* (100, 99.4), *van-B* (100, 98.9), and *vanX-B* (100, 96.7), which collectively is the vanB operon that encodes for vancomycin resistance in *Enterococcus faecium*; while the vanB operon primarily encodes for antibiotic resistance in *E. faecium*, Stinear et al., (^29^2001) have previously isolated vanB-positive anaerobic commensal bacteria from human faeces. We hence reconstructed near-full length 16S rRNA genes to assign taxonomy of operational taxonomic units in our metagenome.

**Table 1:**
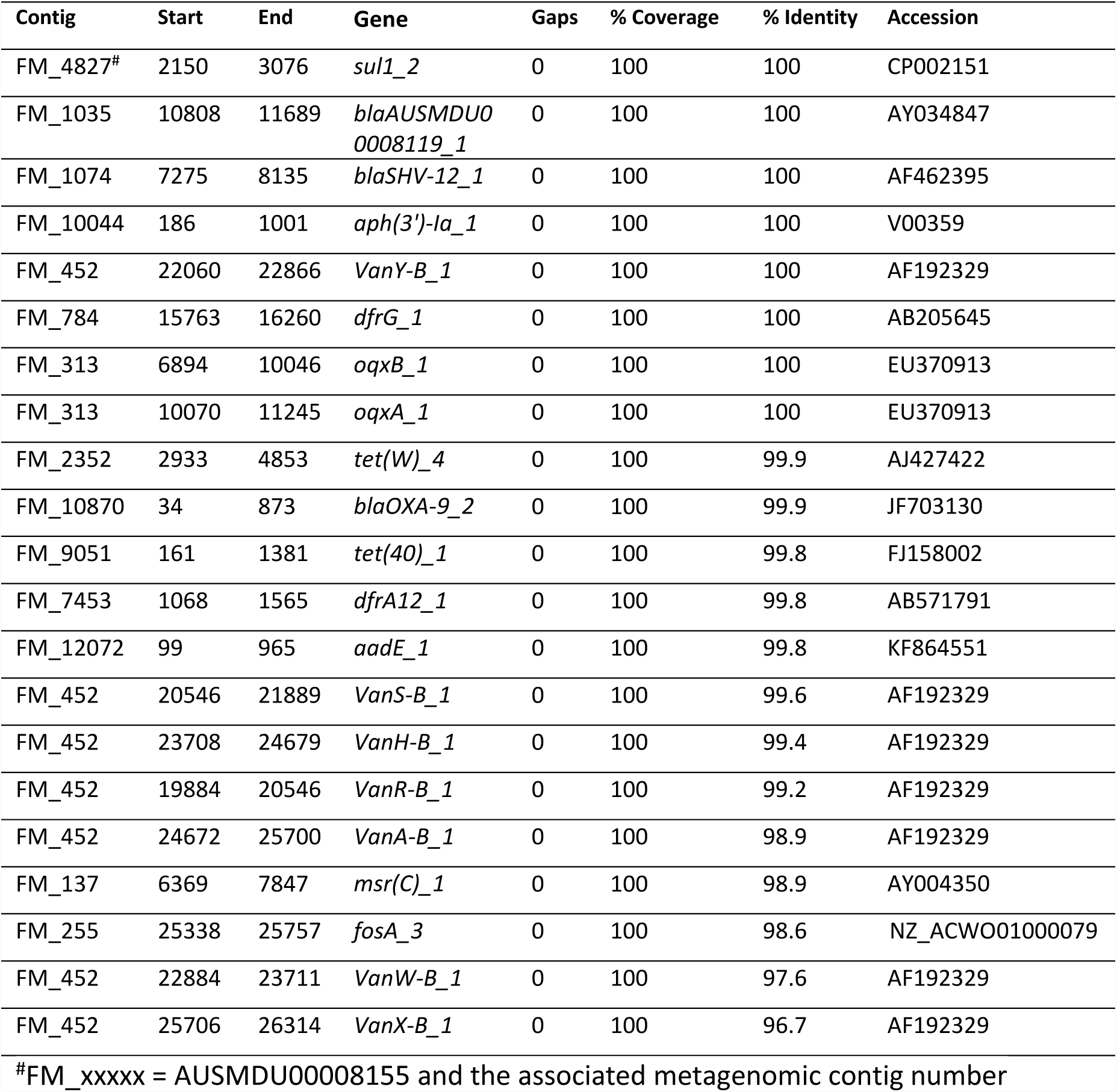
Detection of antimicrobial resistance genes in AUSMDU00008155 metagenome using ABRicate

### Taxonomy of metagenomic reads

Near-full length 16S rRNA genes were reconstructed from the gut microbial community of AUSMDU00008155 to detect the presence of *K. pneumoniae* and *E. faecium*. The Expectation Maximization Iterative Reconstruction of Genes from the Environment program reconstructed fifteen 16S rRNA genes, in which a *K. pneumoniae* 16S rRNA gene was recovered with 100% nucleotide identity over 1161 bp (Table 2). Notably, given the presence of the *vanB* operon, a near-complete 16S rRNA gene at 1344 bp was reconstructed and classified as *E. faecium* at 100% nucleotide identity. Three uncultured organisms were identified as belonging to *Bacteroitdetes*/*Bacteroides*, *Firmicutes*/*Clostridium* XIVa, and *Proteobacteria*/*Sutterellaceae*, with 100% nucleotide identity, and over 1092 bp reconstructed. An independent k-mer based approach (*i.e.,* Kraken) confirmed the presence of *K. pneumoniae*, and *E. faecium* in AUSMDU00008155 metagenomic data.

**Table 2:**
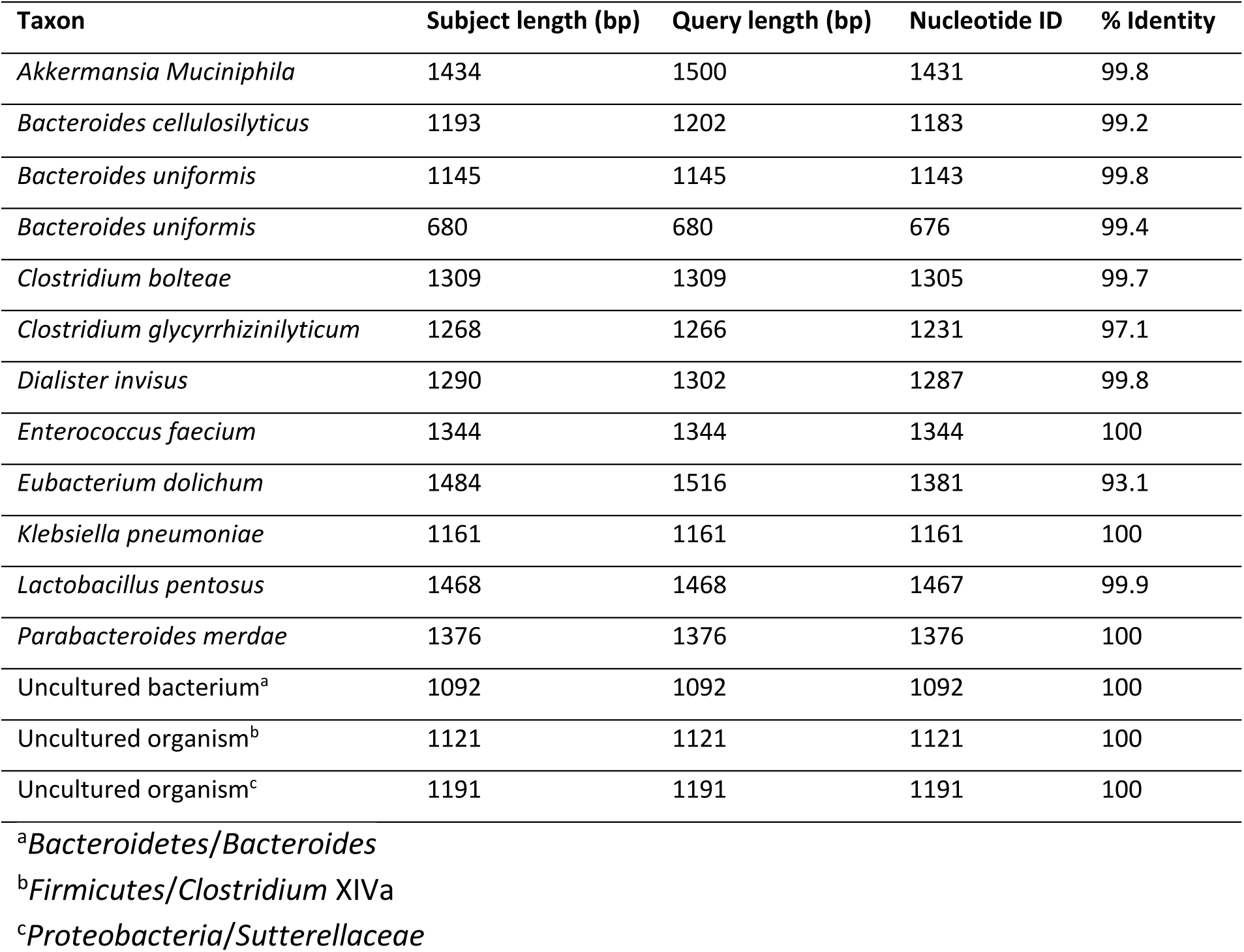
Reconstructed 16S rRNA genes from AUSMDU00008155 metagenome using EMIRGE

| Taxon | Subject length (bp) | Query length (bp) | Nucleotide ID | % Identity |
| --- | --- | --- | --- | --- |
| *Akkermansia Muciniphila* | 1434 | 1500 | 1431 | 99.8 |
| *Bacteroides cellulosilyticus* | 1193 | 1202 | 1183 | 99.2 |
| *Bacteroides uniformis* | 1145 | 1145 | 1143 | 99.8 |
| *Bacteroides uniformis* | 680 | 680 | 676 | 99.4 |
| *Clostridium bolteae* | 1309 | 1309 | 1305 | 99.7 |
| *Clostridium glycyrrhizinilyticum* | 1268 | 1266 | 1231 | 97.1 |
| *Dialister invisus* | 1290 | 1302 | 1287 | 99.8 |
| *Enterococcus faecium* | 1344 | 1344 | 1344 | 100 |
| *Eubacterium dolichum* | 1484 | 1516 | 1381 | 93.1 |
| *Klebsiella pneumoniae* | 1161 | 1161 | 1161 | 100 |
| *Lactobacillus pentosus* | 1468 | 1468 | 1467 | 99.9 |
| *Parabacteroides merdae* | 1376 | 1376 | 1376 | 100 |
| Uncultured bacterium<sup>a</sup> | 1092 | 1092 | 1092 | 100 |
| Uncultured organism<sup>b</sup> | 1121 | 1121 | 1121 | 100 |
| Uncultured organism<sup>c</sup> | 1191 | 1191 | 1191 | 100 |

<sup>a</sup>*Bacteroidetes/Bacteroides*
<sup>b</sup>*Firmicutes/Clostridium XIVa*
<sup>c</sup>*Proteobacteria/Sutterellaceae*

### Reconstruction of isolate genomes from metagenomic reads

Emergent Self Organizing Maps of the tetratnucleotide frequencies of AUSMDU00008155-derived metagenomic contigs reconstructed discrete genome “bins” of isolates from the gut microbial community (Figure 1A). Each point projected onto the ESOM represents DNA fragments 2-5 kb in length, and colour coded with the following convention: AUSMDU00008155 microbiome in red, AUSMDU00008118 in teal, AUSMDU00008119 in navy, and an *E. faecium* AUS0085 strain in purple. A distinct *E. faecium* bin, and a largely mixed bin consisting of closely related AUSMDU00008118 and AUSMDU00008119 derived contigs were resolved (Figure 1B). Furthermore, a “satellite” cluster to the *K. pneumoniae* bin consisted of only AUSMDU00008155 and AUSMDU00008119 contigs, and is typically indicative of mobile genetic elements, such as, plasmids (Figure 1B, *circled*).

**Figure 1.**
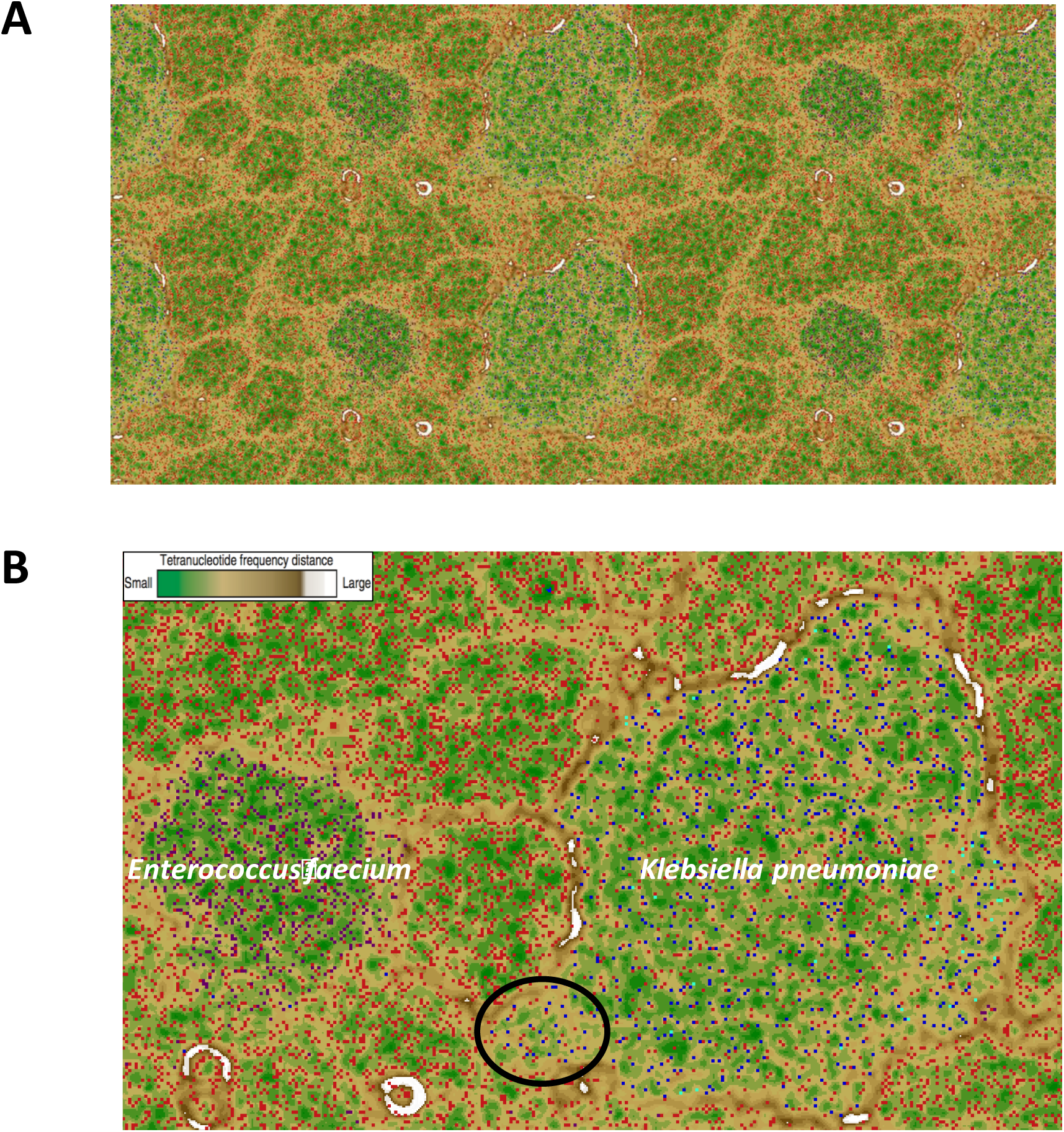
(A) Emergent Self Organizing Map of the tetranucleotide frequencies of AUSMDU00008155 contigs in binned genomes representing (B) *Klebsiella pneumoniae* and *Enterococcus faecium*. Tetranucleotide frequency profiles of DNA fragments between 2 kb to 5 kb in length are projected onto the ESOM. A green background indicates small tetranucleotide frequency distances, white background represents large tetranucleotide frequency distances, while contigs derived from AUSMDU00008155 microbiome is highlighted in red, AUSMDU00008118 in teal, AUSMDU00008119 in navy, and a *vanB*-carrying *E. faecium* in purple.

### Inference of transmission

Using a custom Kraken database, we determined that Patient A’s colonising ST258 *K. pneumoniae* population were most closely related to the reference genome from transmission cluster 2 (AUSMDU00008119), suggesting Patient A was most likely linked to this transmission network (Figure 2). Of the local reference genomes (Kwong *et al., in prep*), AUSMDU00008119 had 136 reads assigned, compared to 11 and 9 reads for the other cluster references. This was corroborated by phylogenetic analysis of the multiple individual colony sequences derived from Patient A’s sample (Kwong *et al., in prep*). These data demonstrate the potential of clinical metagenomics to guide source tracking and infection control efforts.

**Figure 2:**
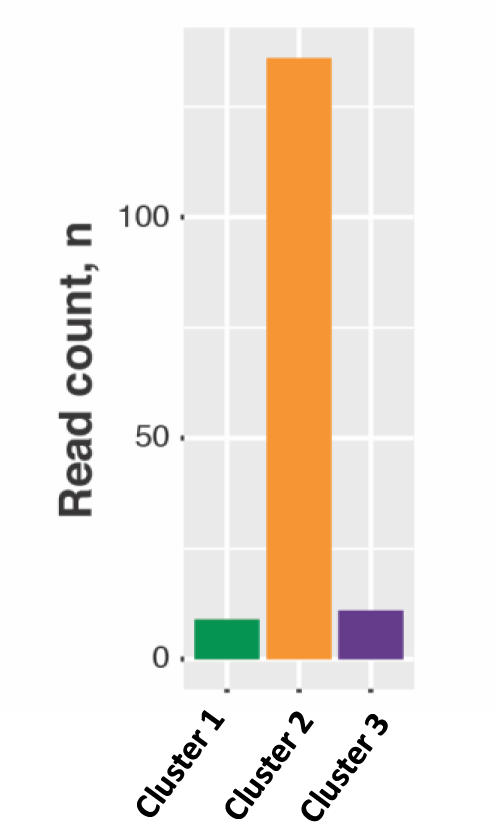
Metagenomic attribution of transmission clusters. The bar graph shows the number of metagenomic reads assigned to each of the reference genomes, representing different transmission clusters described in Kwong *et al.,* (*in prep*).

## Discussion

Previous clinical metagenomic analyses have focused on understanding gut microbiota community diversity 16S rRNA gene level, or incorporating *in vitro* molecular diagnostics as the primary analysis to identify pathogenic isolates ^12^,^30^^-^^32^. In contrast, in this proof-of-principle clinical study, we employed tetranucleotide frequency profiling as the main strategy to reconstruct genomes of community members directly from a faecal specimen, and determine the presence of KPC with strain-level resolution, as well as identifying previously unrecognized colonization with *vanB* VRE. Tetranucleotide profiles (i.e., frequencies of the 256 combinations of G,A,T, and C, in each contig) are a fundamental characteristic of DNA. Contigs with similar tetranucleotide profiles are derived from the same isolate, and therefore, projection of the frequency profiles of metagenomic contigs onto ESOMs can reconstruct isolate genomes independently of reference-based alignment approaches, such as BLAST and/or BWA MEM; this highlights the potential to allow strain level molecular characterization even when a reference isolate is not available *a priori* (c.f., ^33^^-^^34^. The fact that this was achievable using a faeces sample further speaks to the validity of this approach, given the microbial complexity of this sample type.

Metagenomics permitted the comprehensive sampling of genomic content from an adult gut microbial community associated with KPC infection. We evaluated the AMR profile, and detected *bla_KPC_* at 100% coverage with zero gaps, and 100% nucleotide identity (Table 1) which encodes for carbapenem resistance in *Klebsiella pneumoniae*. Furthermore, the resolution of our analyses detected the presence of the genes encoding for an entire *vanB* operon, which confers vancomycin resistance in *E. faecium* isolates, at 100% coverage with zero gaps, and greater than 96% nucleotide identity (Table 1). Reconstruction of near full length 16S rRNA genes from metagenomic short read data recovered an 1161 bp 16S rRNA gene classified as *K. pneumoniae,* and 1344 bp 16S rRNA gene belonging to an *E. faecium* isolate, with 100% nucleotide identity and coverage (Table 2). Our metagenomic analyses uncovered VRE colonization in AUSMDU00008155, and initiated culture analysis of the faecal sample for the presence of VRE using selective media, and a retrospective report notifying the appropriate health care institution of a potential VRE carrier; a reporting that would otherwise have been missed by routine diagnostic analyses. Asymptomatic VRE carriage in AUSMDU00008155 may be facilitated by the co-occurrence of *C. bolteae* in the microbiome, which is described to confer protection against VRE by Carballero *et al.,* (^30^2017); this underscores the important roles microbial community members play in regulating infection and immunity, and suggests whole community metagenomics could become a core component of clinical microbiology.

Tetranucleotide frequency-ESOM analysis recovered metagenomic bins representing single isolates from the AUSMDU00008155 gut microbiome (Figure 1A). Encouragingly, we found that the number of reconstructed genomes (i.e., “bins”) correlated with the number of 16S rRNA genes recovered by our EMIRGE analysis. As a way of further validation, reference *E. faecium* (Aus0085), and AUSMDU00008118 and AUSMDU00008119 genomes were included to guide analysis of ESOMs. Figure 1B illustrates a distinct vanB-carrying *E. faecium* bin, and a large *K. pneumoniae* bin with an associated “satellite” cluster. The main *K. pneumoniae* bin consisted of both AUSMDU00008118 and AUSMDU00008119 derived contigs, while the satellite cluster represents unique genomic content (most probably mobile genetic elements; c.f., ^16^) from AUSMDU00008119, and is therefore indicative of strain-level discrimination. Although we have only assessed one faecal specimen in this study, the richness of microbial characterization obtained from this sample using an alignment-free approach shows the potential for applications in clinical and diagnostic microbiology. This potential is particularly evident for clonal pathogens such as those examined here, where MLST results were unable to distinguish between AUSMDU00008118, AUSMDU00008119, and metagenomic reads (i.e., all isolates were ST258). For example, we accurately assigned Patient A’s colonising metagenomic *K. pneumoniae* isolate to a specific hospital infection cluster (Kwong *et al., in prep*). Our ability to rapidly assign patient metagenomic isolates to pre-existing transmission clusters identified during outbreak situations will better inform prevention control measures to limit the spread of extensively drug-resistant pathogens across our hospital network.

In summary, the current study presents a clinical, and primarily culture-independent investigation framework for the genomic profiling of patients colonised with multidrug-resistant pathogens. Analysis of the whole community metagenome sampled from an adult faecal sample revealed the presence of AMR genes conferring resistance to carbapenem and vancomycin, and identification of pathogenic isolates. We have shown that metagenomic binning, using tetranucleotide frequency profiles, can obtain strain-level resolution. A key implication of incorporating metagenomics into routine clinical microbiology includes higher resolution in AMR and pathogen detection, especially the detection of asymptomatic carriage of antibiotic resistant microbes.

## Acknowledgements

The Microbiological Diagnostic Unit Public Health Laboratory is funded by the Victorian Government. The Doherty Applied Microbial Genomics Program is funded by the Department of Microbiology and Immunology at the Peter Doherty Institute for Infection and Immunity, University of Melbourne. This research was supported by grants from the Austin Medical Research Foundation and the CASS Foundation (Ref No. 7113). JCK (GNT1074824), TPS (GNT1008549) and BPH (GNT1105905) are supported by the National Health and Medical Research Council of Australia.

